# EASI-tag enables accurate multiplexed and interference-free MS2-based proteome quantification

**DOI:** 10.1101/225649

**Authors:** Sebastian Virreira Winter, Florian Meier, Christoph Wichmann, Juergen Cox, Matthias Mann, Felix Meissner

**Affiliations:** Department of Proteomics and Signal Transduction; Max Planck Institute of Biochemistry; *82152 Martinsried*, Germany; Systems Biochemistry; Max Planck Institute of Biochemistry; 82152 Martinsried, Germany; Experimental Systems Immunology; Max Planck Institute of Biochemistry; 82152 Martinsried, Germany

## Abstract

We developed EASI-tag (<u>E</u>asily <u>A</u>bstractable <u>S</u>ulfoxide-based <u>I</u>sobaric tag), a new generation of amine-derivatizing and sulfoxide-containing isobaric labelling reagents, which dissociate at low collision energy and generate peptide-coupled, interference-free reporter ions with high yield. Efficient isolation of ^12^C precursors and quantification at the MS2 level enable accurate determination of quantitative differences between multiplexed samples. EASI-tag makes isobaric labeling applicable to any bottom up proteomics workflow and to benchtop mass spectrometers.

---

Mass spectrometry (MS)-based proteomics has matured remarkably and is now applied in a wide variety of areas^1–3^. It delivers increasingly comprehensive identifications of the constituents of biological systems^4^, however, their accurate quantification across multiple conditions remains an area of active development^5,6^. While label-free quantification is universally applicable and has become widely used due to advances in data acquisition schemes and algorithms, it is limited in sample throughput and achievable quantification accuracy. Isotopic labeling methods allow multiplexing and provide relative quantitation in the same spectra at either the MS1 (full spectrum) or MS2 (fragment spectrum) level. In particular, isobaric labeling methods - such as TMT^7^ or iTRAQ^8^ – have become popular because digested proteins from any sample can be quantified. They generate low-molecular mass reporter ions, in which the intensity of each of the reporters derives from the corresponding isotope-labeled peptide. The major drawback of these methods is the fact that isolation of the precursor ions inevitably co-isolates other precursors, distorting the reporter patterns. This ‘ratio compression’ effect has been addressed with narrow isolation windows, software corrections, additional gas phase manipulation or further fragmentation of the peptide fragment ions^9–12^. However, these methods still do not achieve accurate quantification, or suffer from the decreased sensitivity and acquisition speed caused by their complexity. The fragmentation event leading to low mass reporter ions can also generate precursor ions with the remnant of the tag^13^ and this has been used for quantification^14,15^. Although promising, this method is limited by inefficient generation of the peptide-couple reporters and the fact that the isotope distributions of different channels overlap.

To broaden the applicability of isobaric labeling and to enable determination of accurate ratios on benchtop mass spectrometers, we set out to develop a novel molecule that would generate peptide-coupled reporters with a high yield. Inspired by readily MS-cleavable crosslinking reagents^16^, we devised a sulfoxide-based moiety that dissociates at collision energies below those required for peptide backbone fragmentation. We modified the symmetric to an asymmetric sulfoxide cleavage site, so that fragmentation generates only one instead of three different ion species for each tagged peptide (**Supplementary Fig. 1**). This decreases spectral complexity and increases individual fragment ion intensities, improving sensitivity and computational interpretation of fragmentation spectra. Incorporation of a primary amine reactive moiety for covalent coupling of peptides via an active succinimide ester makes the reagent compatible with standard labeling procedures (**Fig. 1a**). The molecule, which we term EASI-tag - Easily Abstractable Sulfoxide-based Isobaric tag, can be isotope labeled for multiplex quantification and was synthesized in a triplex version for this study. In contrast to other isobaric tags, fragmentation of EASI-tag-labeled peptides produces a neutral loss, thereby retaining the charge state of the precursor (**Fig. 1b**).

**Figure 1.**
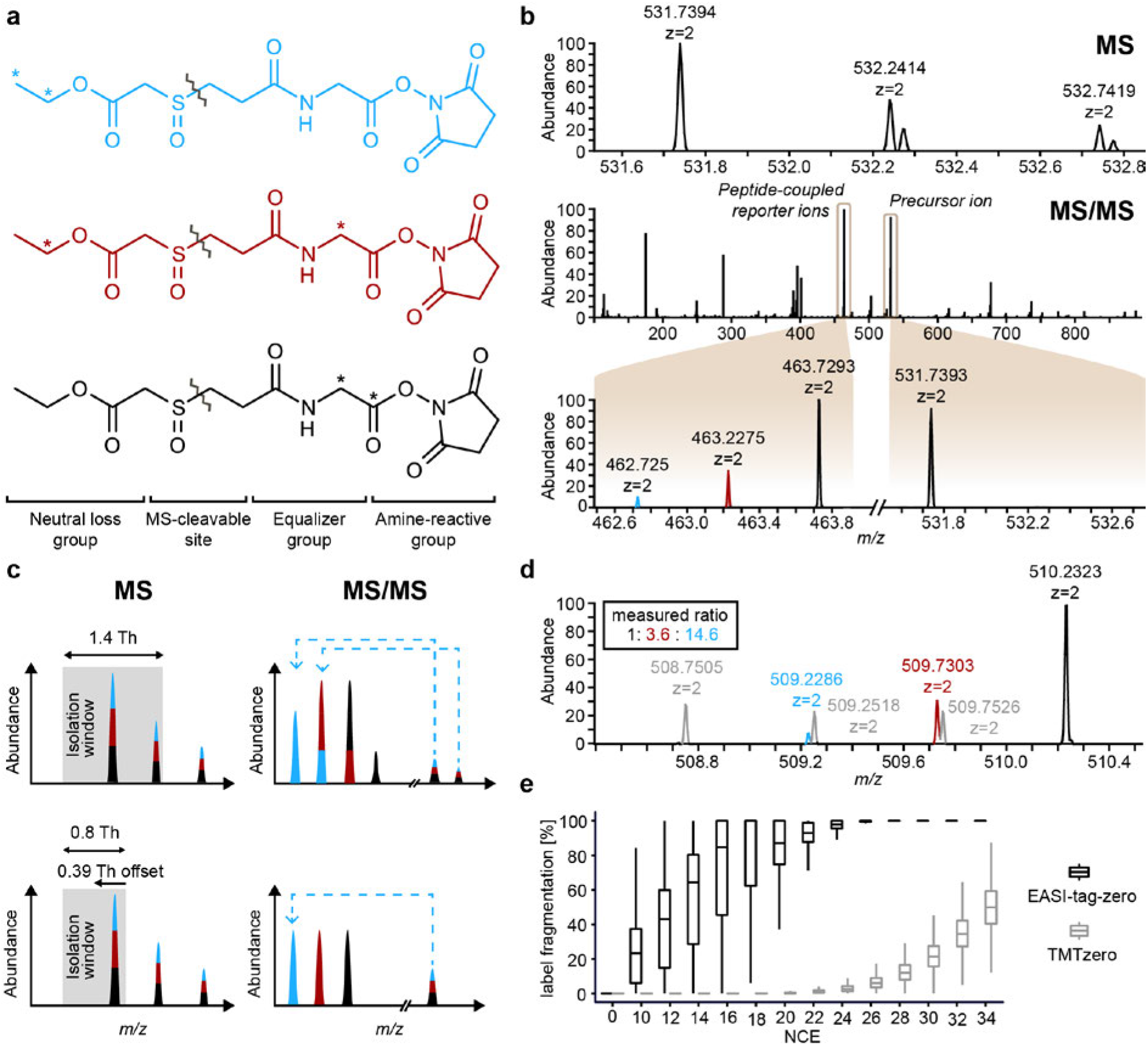
Design and concept of accurate and interference-free MS2-based proteome quantification with EASI-tag. (**a**) Molecular structures of the triplex version of EASI-tag. The isobaric labeling reagents are composed in a modular way of four functional groups and feature a central sulfoxide moiety, which introduces an asymmetric, low-energy cleavage site (zig-zag lines indicate fragmentation site). The stable-isotope labeled positions of the neutral loss and equalizer group for multiplexing are indicated by asterisks. Standard labeling protocols can be applied to couple peptides via the amine-reactive moiety. (**b**) Mass spectra of an EASI-tag-labeled yeast peptide mixed in a ratio of 1:3:10. HCD fragmentation of the doubly charged precursor ion abstracts the neutral loss group and yields the peptide-coupled reporter ion cluster. (**c**) Co-isolation of the natural isotope cluster in a standard isolation window centered on the precursor ion (upper panel) convolutes the relative abundance of peptide-coupled reporter ions. An asymmetric isolation window (lower panel) that suppresses the signal from adjacent isotope peaks and enables direct quantification of reporter ions. (**d**) The precursor mass information is retained in the peptide-coupled reporter ions for EASI-tag labeled peptides. Colored peaks indicate the peptide-coupled reporter ions from an identified yeast peptide in a two proteome experiment (mixing ratios: 1:3:10 for yeast & 1:1:1 for human). Grey peaks are peptide-coupled reporter ions from a co-isolated peptide. (**e**) EASI-tag- and TMT-labeled HeLa peptides were fragmented with normalized collision energies between 10 and 34. (N = 17,565 precursors for EASI-tag & 20,610 for TMT)

Isolation of the entire natural isotope cluster for MS/MS leads to interference of the ^13^C peak of each channel with the ^12^C peak of the adjacent channel, complicating quantification^14^. To address this, and still assure adequate transmission of the precursor, we devised an asymmetric isolation window around the ^12^C isotope, with an optimized transmission width of +- 0.4 Th and an offset of – 0.39 Th (**Fig. 1c** and **Methods**). This effectively suppressed the ^13^C peak to a mean of 1% for doubly and 2% for triply charged precursors, which can be corrected computationally (**Supplementary Fig. 2**). Note that co-eluting peptides that are very close in mass can be quantified separately due to the high resolution employed in the EASI-tag experiment (**Fig. 1d**).

We investigated the fragmentation behavior of thousands of EASI-tag-coupled peptides from a HeLa digest, by varying the normalized collision energies (NCEs) for each of them (**Methods**). This confirmed that - compared to TMT - EASI-tag fragments efficiently at lower collision energies (**Fig. 1e**), and with median NCEs below those required for peptide backbone fragmentation (**Supplementary Fig. 3**). Conceptually, this opens up the possibility for separately optimizing for peptide quantification and identification, which we implemented in the experiments shown here (**Methods**). In Orbitrap mass spectrometers, fragmentation products from these two collision energies can be combined and read out in a single spectrum.

With the EASI-tag and a tailored acquisition strategy in place, we evaluated complex proteome quantification using the isobaric triplex version of our molecule (**Fig. 1a**). We labelled tryptic yeast peptides with the EASI-tag, mixed them 1:3:10 with a 1:1:1 background of human peptides from the HeLa cancer cell line according to a two-proteome model^11^ and analyzed the mixtures by single run LC-MS/MS on an Orbitrap HF instrument (**Methods**).

We implemented the analysis of EASI-tagged peptides in MaxQuant^17^ resulting in 70–77% quantifiable MS2 scans in single runs (**Fig. 2a**). The peptide ratio distributions centered on the values expected from the mixing ratios and were equal in the two and one proteome experiment, indicating the absence of any compression effects (**Fig. 2b**). It is very difficult to measure large ratios in isobaric labeling experiments, particularly when performed at the MS2 level^6,18^. Encouraged by our results, we next tested very large ratios by mixing digested yeast proteomes in 1:12:144 proportions. Again, the measured ratios reflected the mixing ratios, without any evidence of compression (**Fig. 2c**). This capability could be crucial in biological situations where very large fold-changes are expected, such as in the regulated secretome^19^, or when admixing a reference channel in a quantity that makes it likely to be picked for sequencing in the MS1 spectrum.

**Figure 2.**
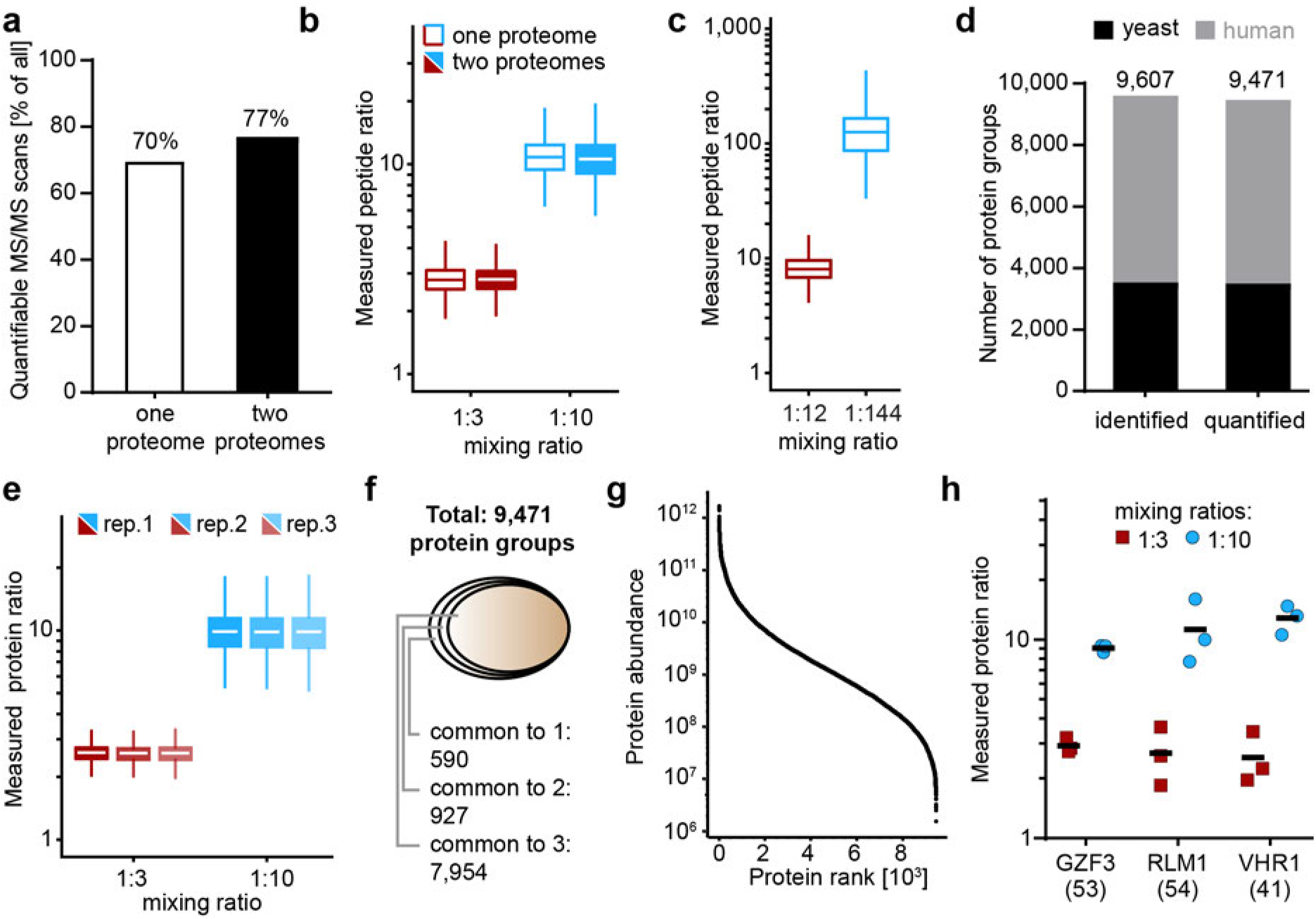
Proteome-wide evaluation of multiplexed quantification with EASI-tag. (**a**) MS/MS scans in the one proteome/two proteomes experiments with intensity signals in at least two peptide-coupled reporter channels. (**b**) Quantified peptide-coupled reporter ion ratios (log10) of yeast peptides mixed in a 1:3:10 ratio in the absence or presence of a HeLa background proteome (mixed 1:1:1) in a single run LC-MS/MS experiment (N=7,419 scans & 3,055 scans for the 1:3 mixing ratio and N=6,729 scans & 2,622 scans for the 1:10 mixing ratio). (**c**) Quantified peptide-coupled reporter ion ratios (*log*10) of yeast peptides mixed in a 1:12:144 ratio in a single run LC-MS/MS experiment (N=6,606 scans for the 1:12 mixing ratio and N=2,642 scans for the 1:144 mixing ratio). (**d**) A total of 9,607 protein groups were identified in 16 high pH fractionations of the mixed yeast/HeLa two proteome model in triplicate LC-MS/MS runs. Out of these, 9,471 protein groups were quantified with intensity signals in at least two peptide-coupled reporter channels. (**e**) Ratio distribution (*log*10) of quantified yeast protein groups from (**d**) (N=3,160, 3,212, 3,139 for the 1:3 mixing ratio and N=3,022, 3,059, 2,990 for the 1:10 mixing ratio). (**f**) 92% (8,881) of the identified protein groups were quantified in at least two replicate experiments. (**g**) Ranked abundance of yeast and human protein groups quantified with EASI-tag. (**h**) Accurate quantification of low abundant proteins such as yeast transcription factors with EASI-tag. Estimated copy numbers from Kulak et al.^20^ are indicated in parenthesis.

To investigate the EASI-tag method in global proteome analysis, we fractionated the two-proteome experiment into 16 fractions and analyzed each fraction in triplicate 100 min gradients. Overall, accurate quantification was obtained from more than 300,000 MS/MS scans (**Supplementary Fig. 4a**). We identified 9,607 protein groups (3,535 yeast) with a mean number of 8.5 peptides (**Fig. 2d**). Even at this depth, the ratios closely matched the expected values and 92% of all identified protein groups were quantified in at least two of the triplicates (**Fig. 2e, f**). Quantification accuracy of these proteins reached a median CVs of 6.7% and 12.2% for the two reporter ion channel ratios over the entire abundance range (7.3% and 16.2% for the 1:3 and 1:10 yeast mixing ratios) (**Supplementary Fig. 4b**). EASI-tag based proteome quantification covers an abundance range over six orders of magnitude (**Fig. 2g**) and even yeast transcription factors present at less than 100 copies per cell^20^ were quantified correctly (**Fig. 2h**).

The EASI-tag strategy has all the advantages of existing isobaric labeling methods but adds important new dimensions. The absence of ratio compression assures that accurate as well as precise fold-changes can be measured. Such a capability is desired in many biomedical areas, for instance the absolute quantification of expression levels or concentrations in clinical measurements where spiked in reference standards are required. While we established the concept of accurate and interference-free MS2-based proteome quantification with the triplex version of EASI-tag, its multiplexing capability can be readily extended in the future. In combination with the streamlined analysis workflow integrated in MaxQuant, our strategy is universally applicable on benchtop instruments and paves the way for wide-spread use of isobaric labeling in a large number of laboratories.

## Methods

### Labeling reagents

TMTzero was purchased from Thermo Fisher Scientific. The EASI-tag labeling technology was conceptualized by the authors and custom chemical synthesis of the EASI-tag reagents was performed by Taros Chemicals GmbH, Germany. For proof-of-principle experiments, a non-isotopically coded version (‘EASI-tag-zero’), and for MS2-based proteome quantification, three isotopomers (‘triplex EASI-tag’) containing two ^13^C labeled carbons at the positions indicated in Figure 1a were used.

### Cell culture

The human cervical cancer cell line (HeLa S3 clone, ATCC) was cultured in Dulbecco’s modified Eagle’s medium, supplemented with 10% fetal bovine serum, 20 mM glutamine and 1% penicillin-streptomycin (all PAA Laboratories). The cells were harvested by centrifugation and washed once with cold phosphate buffered saline before re-centrifugation. Saccharomyces cerevisiae strain BY4741 (EUROSCARF) was grown to log phase at 30°C in yeast extract peptone dextrose medium (10 g/L BactoYeast extract, 20 g/L peptone and 2% w/v glucose) and harvested by centrifugation. The cells were washed with cold Milli-Q water and pelleted by centrifugation. All cell pellets were flash frozen in liquid nitrogen and stored at −80°C.

### Lysis and digestion

Cell lysis as well as protein reduction and alkylation were performed in a one-pot SDC buffer system containing chloroacetamide^20^ (PreOmics GmbH). Briefly, HeLa and yeast cell pellets were re-suspended, boiled at 95°C for 10 min and sonicated 15 cycles for 30 s with a Bioruptor Plus sonication device (Diagenode) to enhance cell disruption before adding equal amounts of the proteases LysC and trypsin in a 1:100 (w/w) ratio. The enzymatic digestion was performed overnight and stopped with trifluoroacetic acid (TFA) at a final concentration of 1% (v/v). The acidified protein digests were desalted on C18 StageTips^21^ and eluted with 80% acetonitrile/0.1% TFA (v/v).

### Isobaric labeling

Approximately 20 μg of desalted peptide mixtures were reconstituted in 50 mM HEPES buffer (pH 8.5). The EASI-tag or TMTzero isobaric label was reconstituted in water-free acetonitrile and added in 4-fold excess (w/w) to couple primary and N-terminal amino groups to the N-Hydroxdysuccinimide (NHS)-activated ester. Acetonitrile was added to a final concentration of 33% (v/v) and peptides were labeled for 1 hour at 25 °C. Unreacted NHS ester was hydrolyzed by adding 10 volumes of 1% TFA. The labeled peptides were mixed at defined ratios, purified on C18 StageTips with 2% acetonitrile/0.1% TFA and eluted as above. The eluate was evaporated to dryness in a vacuum centrifuge and the peptides were re-constituted in 2% acetonitrile/0.1% TFA for single run LC-MS/MS analysis or high-pH reverse phase fractionation.

### High pH reversed-phase fractionation

Purified protein digests were fractionated using a 30 cm reversed-phase column (I.D. 250 um) with a ‘spider fractionator’^22^ (PreOmics GmbH) system coupled online to an EASY-nLC 1000 chromatograph (Thermo Fisher). For EASI-tag-labeled peptides, separation was performed at a flow rate of 2 uL/min with a binary buffer at pH 10 (PreOmics GmbH) starting from 10% B, followed by a stepwise increase to 45% B within 75 min and 95% B within 7 min. For TMT-labeled peptides, the gradient started from 3% B, followed by stepwise increases to 30% B within 45 min, to 40% B within 12 min, to 60% B within 5 min and finally to 95% B within 10 min.

In the deep two-proteome experiment, sixteen fractions were automatically concatenated by shifting the rotor valve every 120 s and collecting into 0.2 mL tubes. To collect eight fractions (for breakdown curves, see below), the rotor valve was shifted every 100 s. Collected fractions were vacuum-centrifuged to dryness and re-constituted in 2% acetonitrile/0.1% TFA for LC-MS/MS analysis.

### Liquid chromatography and mass spectrometry

LC-MS/MS analysis was performed on an EASY-nLC 1200 ultra-high pressure system coupled online to a Q Exactive HF mass spectrometer via a nano-electrospray ion source (all Thermo Fisher Scientific). Approximately 2 μg of peptides were separated at a constant flow rate of 300 nL/min at 60 °C on a 45 cm reverse-phase column (I.D. 75 μm) packed with ReproSilPur C18-AQ 1.9 μm resin (Dr. Maisch GmbH). Mobile phases A and B were 100/0.1% water/formic acid (v/v) and 80/20/0.1% acetonitrile/water/formic acid (v/v/v). For EASI-tag labeled samples, the initial concentration of 10% B was linearly increased to 45% B within 95 min, followed by further increases to 60% within 5 min and 95% B within 5 min and a 5 min plateau before re-equilibration. TMTzero-labeled and unlabeled peptides were separated as above but with initial gradients from 10 to 35% B and from 5 to 30% B, respectively.

Proteomics experiments were acquired with a data-dependent top10 method. Full MS scans were acquired from *m/z* 300-1,650 at a resolution of 60,000 at *m/z* 200. The AGC target was set to 3e6. Higher energy collisional dissociation MS/MS scans were acquired with a stepped normalized collision energy of 11 and 21 (50% each) at a resolution of 60,000 at *m/z* 200. Precursor ions were isolated in an asymmetric 0.8 Th window (offset −0.39 Th) and accumulated to reach an AGC target value of 2e5 or for a maximum of 110 ms. Only doubly and triply charged precursors were selected for fragmentation and the monoisotopic peak was preferably isolated. The acquisition software was adapted via the Thermo Fisher application programming interface (API) to implement the latter functionality. Precursors were dynamically excluded for 30 s after the first fragmentation event.

To investigate the fragmentation behavior of EASI-tag-zero- and TMTzero-labeled peptides, we acquired data-dependent breakdown curves with ‘Full MS’ parameters set as above. Higher energy collisional dissociation MS/MS scans were acquired at a resolution of 30,000 at *m/z* 200. AGC target value and maximum ion injection time were set to 2e5 and 60 ms, respectively. Suitable precursor ions were repeatedly isolated with a symmetric isolation window of 1.5 Th fourteen times per acquisition cycle with increasing collision energy. The normalized collision energy was set to 0 for the first scan and subsequently increased from 10 to 34 with a step size of 2.

To evaluate the ion transmission characteristics of the analytical quadrupole, we acquired a similar experiment with an unlabeled HeLa digest. The collision energy was fixed at 0 and the center of the quadrupole isolation window was shifted incrementally to lower *m/z* within one acquisition cycle, starting from the precursor-centered position. In the initial experiment, for three different isolation widths (0.8, 1.0 and 1.4 Th), the isolation offset was 0 for the first scan and increased from −0.3 to −0.8 Th with 0.05 Th increments for the subsequent scans. For a more detailed evaluation, we varied the offset from −0.35 to −0.45 Th with increments of 0.01 Th for an isolation window of 0.8 Th.

### Data analysis

Raw spectral data were extracted with the MSFileReader (v3.0.31, Thermo Fisher Scientific) and proteomics MS Raw files were processed with MaxQuant^17^ (version 1.6.0.9) for identification and quantification. The MS/MS spectra were searched against tryptic peptides with less than 3 missed cleavages derived from human and yeast reference proteomes (Uniprot, accessed 02/2017) as well as a list of potential contaminants with the Andromeda search engine. The search included cysteine carbamidomethylation as fixed modification and methionine oxidation as variable modification. For quantitative proteomics experiments, EASI-tag was defined as a modification of the type ‘isobaric label’ at any lysine residue or the peptide N-terminus with a delta mass of 249.0582 Da (^12^C_7_^13^C_2_H_13_O_5_NS). For analysis of the breakdown curves, EASI-tag-zero and TMTzero were defined as variable modifications at any lysine residue or the peptide N-terminus with delta masses of 247.0514 Da (^12^C_9_H_13_O_5_NS) and 224.1525 Da (^12^C_12_H_20_O_2_N_2_), respectively. To avoid analysis of SIM scans (NCE 0 scans) as full scans by MaxQuant, these were removed before MQ analysis. The minimum peptide length was set to 7 amino acid residues and the maximum peptide mass was limited to 4,600 Da. The maximum allowed initial mass deviation was 4.5 ppm for precursor ions after non-linear recalibration and 10 ppm for fragment ions. The initial filtering criteria for MS/MS scans was set to 20 peaks per 100 Da interval and the ‘second peptide’ option was disabled. Where applicable, MS/MS precursor mass values were corrected for the isolation window offset. Identifications were controlled at a maximum false discovery rate of 1% for both, peptide spectrum match and protein level.

For the quantification of EASI-tag coupled reporter ions, a mass tolerance of 15 ppm was set in MaxQuant. Neutral loss masses were defined as 136.0194 Da (^12^C_4_^13^C_0_H_8_O_3_S), 137.0228 Da (^12^C_3_^13^C_1_H_8_O_3_S) and 138.0261 Da (^12^C_2_^13^C_2_H_8_O_3_S) for the reporter ion channels 0, 1 and 2, respectively.

### Bioinformatic analysis

Bioinformatic analysis was performed in Perseus^23^ and the R statistical computing environment. Decoy database hits were strictly excluded from the analysis as well as potential contaminants and proteins that were identified exclusively by one site modification.

For EASI-tag based protein quantification (Figs. 2e and h), we required reporter intensities in at least two channels and in at least two out of three replicates. Coefficients of variation indicate the variability per protein group in replicate measurements.

For peptide quantifications (Figs. 2b and c), reporter ions that fell outside of a 5 ppm mass tolerance window with respect to the median mass estimated from all three peptide-coupled ions were excluded from the analysis. Occasionally, the software selected the ^13^C peak instead of the ^12^C peak. To exclude these cases from analysis, we discarded spectra with peaks 1 Da below the selected precursor ion if its relative abundance was more than 10% of the precursor ion in the full scan (using R and mzXML format). After filtering, peptide-coupled reporter ion ratios were calculated by dividing the reporter intensity in channel 0 by reporter intensities in channel 1 and channel 2.

To determine the fragmentation efficiencies of EASI-tag-zero and TMTzero, breakdown curves were first processed with MaxQuant for peptide identification. Scan cycles with any MS/MS scan with an AGC fill below 50% were excluded from the analysis. Label fragmentation efficiencies were calculated by dividing the intensity of the peptide-coupled reporter ion by the summed intensities of the precursor ions with intact and fragmented label. Only scan cycles with identified, labeled peptides were included in the analysis and if either the precursor ion or the peptide-coupled reporter ion were not detected, its intensity was set to 0. If both, the precursor and peptide-coupled reporter ion were not detected, the scan was excluded from the calculation.

For Supplementary Figure 3, the NCE of the label fragmentation was defined as the minimal NCE at label fragmentation (see above) reached 25%. If label fragmentation did not reach 25%, the NCE of the label fragmentation was defined as the NCE of the maximal label fragmentation. The NCE of peptide backbone fragmentation was defined as the NCE at which the sum of the precursor and peptide-coupled reporter intensities was below 20% of the precursor intensity in the SIM scan (NCE = 0). If the sum of the precursor and peptide-coupled reporter intensities did not drop below 20% of the precursor intensity in the SIM scan, the NCE of peptide backbone fragmentation was defined as the NCE at which the sum of the precursor and peptide-coupled reporter intensities reached its maximum.

To evaluate the quadrupole transmission, precursor ion intensities and intensities of the M+1 ion in MS/MS scans from doubly or triply charged precursors in the center of the gradient (scan numbers 8,000-40,000) were analyzed. Zero values were removed before mean calculation.

### Data presentation

Boxplot elements of all figures are defined as follows: center line = median, lower and upper box limits = first and third quartiles, whiskers = maximum 1.5x inter-quartile range, outliers not displayed.

## Acknowledgements

We acknowledge all members of the Department of Proteomics and Signal Transduction for help and valuable discussions. We thank Philipp Geyer and Sophia Doll for assistance with the experiments; and Gaby Sowa, Igor Paron and Korbinian Mayr for technical support.

This study war partially supported by funding from the German Research Foundation (DFG/Gottfried Wilhelm Leibniz Prize), European Union’s Horizon 2020 research and innovation program under grant agreement no. 686547 (MSmed project) and the Max-Planck Society for the Advancement of Sciences.

## Author contributions

FeM and MM conceived the study; FeM and SVW conceptualized and designed the EASI-tag; FlM, FeM and MM conceptualized the acquisition method; and CW contributed to its implementation. SVW, FlM and CW performed the experiments; JC developed and implemented data analysis software; SVW, FlM, FeM and MM analyzed the data; SVW, FlM, FeM and MM wrote the manuscript with input from all authors.

